# Surface proteins of SARS-CoV-2 drive airway epithelial cells to induce interferon-dependent inflammation

**DOI:** 10.1101/2020.12.14.422710

**Authors:** Gautam Anand, Alexandra M. Perry, Celeste L. Cummings, Emma St. Raymond, Regina A. Clemens, Ashley L. Steed

## Abstract

SARS-CoV-2, the virus that has caused the COVID-19 pandemic, robustly activates the host immune system in critically ill patients. Understanding how the virus engages the immune system will facilitate the development of needed therapeutic strategies. Here we demonstrate both in vitro and in vivo that the SARS-CoV-2 surface proteins Spike (S) and Envelope (E) activate the key immune signaling interferon (IFN) pathway in both immune and epithelial cells independent of viral infection and replication. These proteins induce reactive oxidative species generation and increases in human and murine specific IFN-responsive cytokines and chemokines, similar to their upregulation in critically ill COVID-19 patients. Induction of IFN signaling is dependent on canonical but discrepant inflammatory signaling mediators as the activation induced by S is dependent on IRF3, TBK1, and MYD88 while that of E is largely MYD88 independent. Furthermore, these viral surface proteins, specifically E, induced peribronchial inflammation and pulmonary vasculitis in a mouse model. Finally we show that the organized inflammatory infiltrates are dependent on type I IFN signaling, specifically in lung epithelial cells. These findings underscore the role of SARS-CoV-2 surface proteins, particularly the understudied E protein, in driving cell specific inflammation and their potential for therapeutic intervention.

**Author Summary:** SARS-CoV-2 robustly activates widespread inflammation, but we do not understand mechanistically how the virus engages the immune system. This knowledge will facilitate the development of critically needed therapeutic strategies to promote beneficial immune responses will dampening harmful inflammation. Here we demonstrate that SARS-CoV-2 surface proteins spike and envelope alone activated innate cell function and the interferon signaling pathway. This activation occurred in both immune and epithelial cells, and mechanistic studies demonstrated dependence on known key inflammatory signaling mediators, IRF3, TBK1, and MYD88. In animal studies, we showed that these viral surface proteins induce epithelial cell IFN-dependent lung pathology, reminiscent to acute COVID-19 pulmonary infection. These findings underscore the need for further investigation into the role of SARS-CoV-2 surface proteins, particularly the understudied E protein, in driving cell specific inflammation.

## Introduction

SARS-CoV-2 infection has profoundly impacted human health globally, leading to more than 70 million cases and 1,600,000 deaths as of December 11, 2020. The ensuing illness, termed COVID-19, predominantly manifests as a respiratory disease, which disproportionately affects the older population and those with comorbidities. Many critically ill patients with COVID-19 develop respiratory failure characterized by poor gas exchange and damaging lung inflammation (1, 2).

This novel virus was quickly identified as a beta-coronavirus that has 79.5% genetic similarity with SARS-CoV (Severe Acute Respiratory Syndrome-Coronavirus) and 50% with MERS (Middle East Respiratory Syndrome) (3-5). SARS-CoV-2 also shares a host receptor with SARS-CoV for cell entry, namely, angiotensin-converting enzyme 2 (ACE2), via the binding of its surface protein Spike (S) (4, 6, 7). The S protein of SARS-CoV-2 binds ACE2 more avidly than that of SARS-CoV, although these two S proteins share similar tertiary structures (8). Genomic comparison of SARS-CoV-2 with SARS-CoV shows there are 27 changes in the amino acid sequence of S, and the majority of these substitutions occur outside of the ACE2 binding domain (9). However, mutations in key S epitopes may contribute to conformational changes that increase ACE2 affinity, influence antigenicity, and/or affect the ability of SARS-CoV-2 to activate immune responses (10).

While the S protein interaction with ACE2 has been the focus of vaccine design, other structural proteins likely play key roles in disease pathogenesis. The coronaviral genomes also encode structural proteins Nucleocapsid (N), Envelope (E), Membrane (9, 11). However, little is known about these structural proteins’ roles in immune activation and pathogenesis. The N protein has been shown to have an immunomodulatory function in SARS-CoV infection (12, 13). Interestingly, the SARS-CoV and SARS-CoV-2 E proteins have no amino acid substitutions. SARS-CoV E is essential for viral morphology, budding, and tropism (14, 15). Importantly, the SARS-CoV E was found to enhance inflammasome activation (16-19). Therefore, the conserved E protein and its engagement of the host immune response could prove to be a potent therapeutic intervention point useful for targeting multiple coronaviruses if its mechanistic actions are clearly understood (20).

During acute infection, COVID-19 patients are in a seemingly hyperinflammatory state with a dysregulated immune response (21). Similar to other RNA-viral infections, the pulmonary disease of COVID-19 is likely a combination of direct viral damage and this hyperactivated host immune response. While lymphopenia has been a consistent finding in COVID-19 (9, 22, 23), many patients also exhibit a cytokine storm which is associated with disease severity and outcome (7-9, 21, 24-27). These patients demonstrate an increase in number of inflammatory monocytes and elevated serum levels of proinflammatory chemokines and cytokines including IL-2, IL-7, IL-10, IL-6, G-CSF, IP-10, MIP-1α, MCP-1 and TNF-α (1, 2, 7, 21, 25, 26, 28-34). While these chemokines and cytokines attract immune cells to mount an antiviral defense, the resulting cytokine storm and cellular infiltration have been implicated in lung cell damage and disease pathogenesis.

Given the key roles of the innate immune response in both viral clearance and disease pathogenesis, understanding how SARS-CoV-2 structural proteins elicit host immunity is necessary for designing optimal therapeutic strategies. Therefore, we sought to investigate the innate immune response to SARS-CoV-2 antigens, independent of viral infectivity and nuclei acid replication. In this report, we demonstrate that the purified structural proteins of SARS-CoV-2 alone activate inflammatory pathways in immune and epithelial cells and induce localized lung pathology dependent on IFN signaling in epithelial cells. These findings implicate the contribution of the viral surface proteins to driving inflammation in a cell type specific manner and highlight their potential for therapeutic intervention.

## Results

### Purified SARS-CoV-2 proteins induce reactive oxygen species generation and proinflammatory chemokine and cytokine production

Increased reactive oxidative species generation has been detected in clinical COVID-19 sputum samples (35), although it is unclear to what extent active infection or inflammatory stimulation contribute to this finding. To examine the role of the SARS-CoV-2 structural proteins in directly (i.e. the absence of infectious virus) activating this innate immune cell effector function, we investigated the ability of S and E antigens to induce ROS generation in macrophages. While SARS-CoV-2 does not infect wildtype (wt) mice *in vivo (36)*, S protein and an N-terminal 10 amino acid truncated envelope protein (E-Trunc) potently enhanced zymosan-induced ROS generation in *ex vivo* isolated wt murine peritoneal macrophages after overnight incubation compared to control samples by 2.09 ± 0.35-fold and 2.63 ± 0.95-fold, respectively. (Fig 1A). Alveolar macrophages also demonstrated increased zymosan-induced ROS production in response to E-Trunc (1.71±0.19-fold) but not in response to S (Fig 1B).

**Figure 1.**
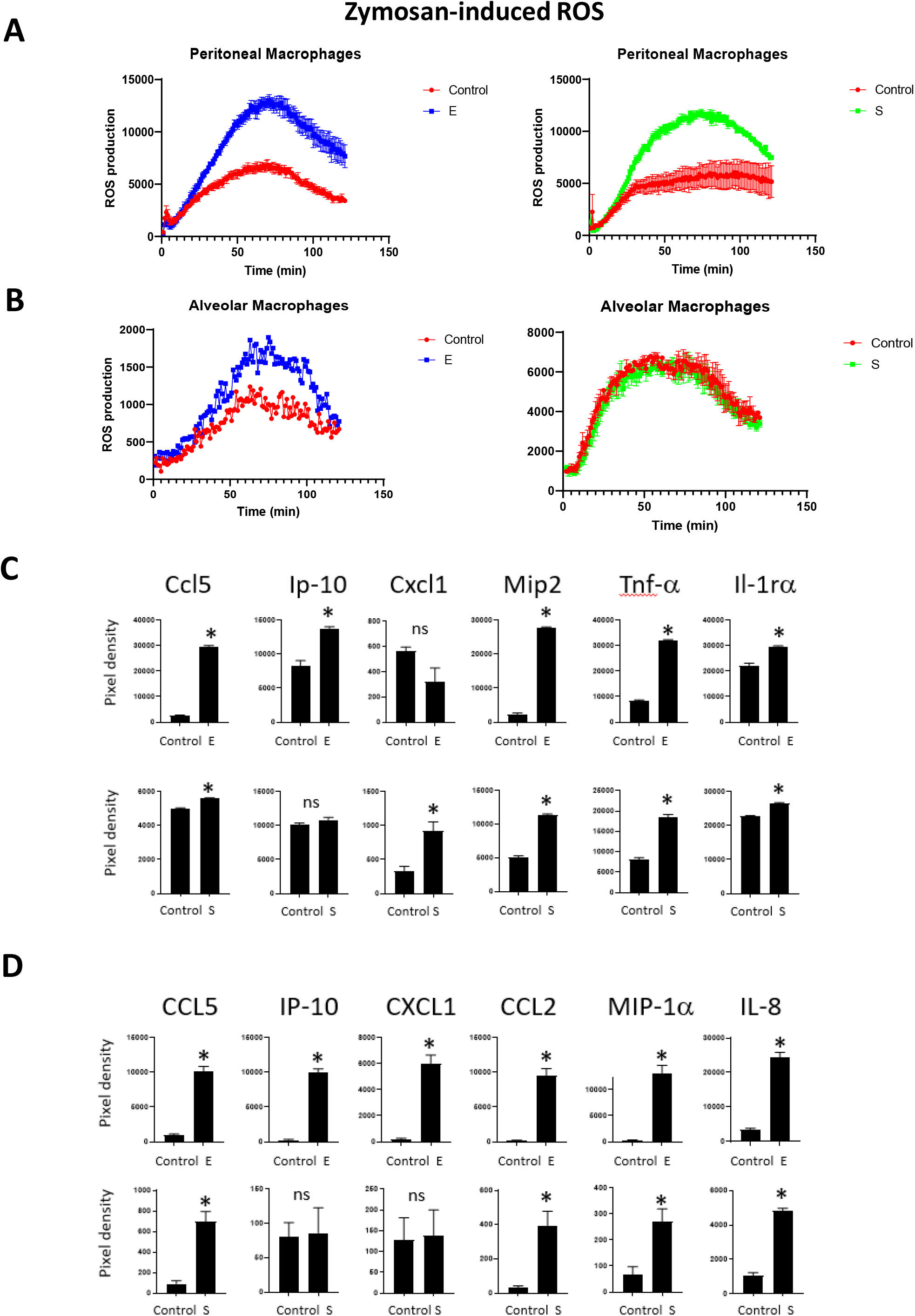
SARS-CoV-2 truncated Protein E and S Enhance ROS Production and Increase the Expression of Proinflammatory Chemokines and Cytokines. (A) ROS production in peritoneal (A) and alveolar macrophages (B) after zymosan (20µg/ml) stimulation and incubation with SARS-CoV-2 peptides E-Trunc and S. (Representative figures with n=2 experiments with 6 mice per experiment). (C-D) Detection of chemokines and cytokines in the culture supernatant of RAW (C) and THP1 cells (D) incubated with E-Trunc or S (2 µg/ml) or control for 24 hours. The graphs show measurements of the pixel density in dot arrays. (n=2 biological samples for each condition with 2 technical replicates of each sample). Graphs depict average with SEM. *p< or =0.05 and ns denotes not statistically significant. Mann-Whitney was used for statistical analysis.

Given the finding that viral surface proteins induced an increase in an innate immune effector function, we also examined the induction of specific chemokines and cytokines in myeloid inflammatory reporter cell lines. Similar to induction of ROS generation in primary cells, E-Trunc and S induced increases in chemokines and cytokines when incubated with murine IFN reporter RAW cells (RAW-Lucia ISG). Both E-Trunc and S enhanced the following chemokines and cytokines: Ccl5, Mip-2, Il-1rα, and G-Csf (Fig 1C and S1A Fig). E-Trunc peptide independently increased the expression of Ip-10, while the S protein increased Cxcl1, GM-Csf, Il-6, and Il-10. This distinct induction of specific chemokines and cytokines indicates that these viral proteins likely induce host inflammatory responses by different mechanisms.

To determine if human myeloid cells similarly responded to the SARS-CoV-2 structural proteins, we next incubated a monocyte THP1 reporter cell line (THP1-Dual) with E-Trunc and S antigens overnight. Indeed, E-Trunc and S induced both shared and distinct increases in many inflammatory mediators in human monocytes (Fig 1D and S2B Fig). E-Trunc dramatically increased the expression of CCL5 (10-fold), IP-10 (41-fold), CXCL1 (30-fold), CCL2 (48-fold), MIP-1α (57-fold) and IL-8 (7-fold). S protein similarly increased the expression of CCL5 (7.6-fold), CCL2 (12.7-fold), MIP-1α (4.2-fold), and IL-8 (4.5-fold), albeit to a lesser magnitude than increased by E-Trunc. CCL5 transcript expression by S and E-Trunc was confirmed by qRT-PCR (S2A Fig). Interestingly, S protein alone specifically increased IL-1Rα (2.3-fold), GM-CSF (2.9-fold), and CXCL12 (1.5-fold). These findings underscore that there are shared as well as distinct immune responses to specific coronavirus surface antigens and implies unique mechanisms of activation.

Increased serum TNF-α has been found during COVID-19 infection (9, 25, 26, 31). Previous work has also shown a specific increase in TNF-α expression in mouse macrophages by the SARS-CoV S protein (37). Likewise, we also found that TNF-α increased in mouse monocytes incubated with SARS-CoV-2 proteins E-Trunc or S (Fig 1C). However in human myeloid cells our results were inconsistent; there was no difference in the levels of TNF-α after exposure to S and a small decrease (0.7-fold) in protein but increased mRNA transcript after incubation with E-Trunc (S1B-S2A Fig). These results highlight key commonalities and differences in inflammatory responses between human and mouse cells. This knowledge bears critical attention as we rely on animal models to investigate SARS-CoV-2 mechanistically and test new therapeutic strategies.

### Purified SARS-CoV-2 proteins induce inflammatory signaling

As our findings above showed that E-Trunc and S protein upregulate multiple chemokines and cytokines known to be IFN responsive, we directly asked whether these antigens activate IFN induction. We incubated the IFN reporter cell lines, which harbor tandem interferon stimulated response elements inducing luciferase expression, with E-Trunc and S as well as the SARS-CoV-2 structural protein N and the full length E protein (E-Full). After 24-hour incubation, IFN induction were enhanced in both murine and human monocytes, most robustly by E-full in the human THP-1 reporter cells and E-Trunc in the murine RAW reporter cells (Fig 2A). E-Full enhanced luciferase expression by 3-fold in RAW cells and 6.5-fold in THP-1 cells while E-Trunc lead to a 5.5-fold and 2.8-fold increase, respectively. Protein S enhanced IFN signaling in these cells to a lesser extent by 1.3-fold in RAW cells and 1.5-fold in THP-1 cells. A VSV-pseudovirus expressing the SARS-CoV-2 S protein on its virion surface also increased IFN induction in both THP-1 and RAW cells compared to VSV expressing its glycoprotein (VSV-G), albeit with observed cytotoxicity in THP-1 VSV-G infected cells (Fig 2B-C). Importantly, the structural protein N did not induce IFN signaling in any of the cell lines tested. Of note, the N protein is contained inside the virion while E and S are displayed on the viral surface.

**Figure 2.**
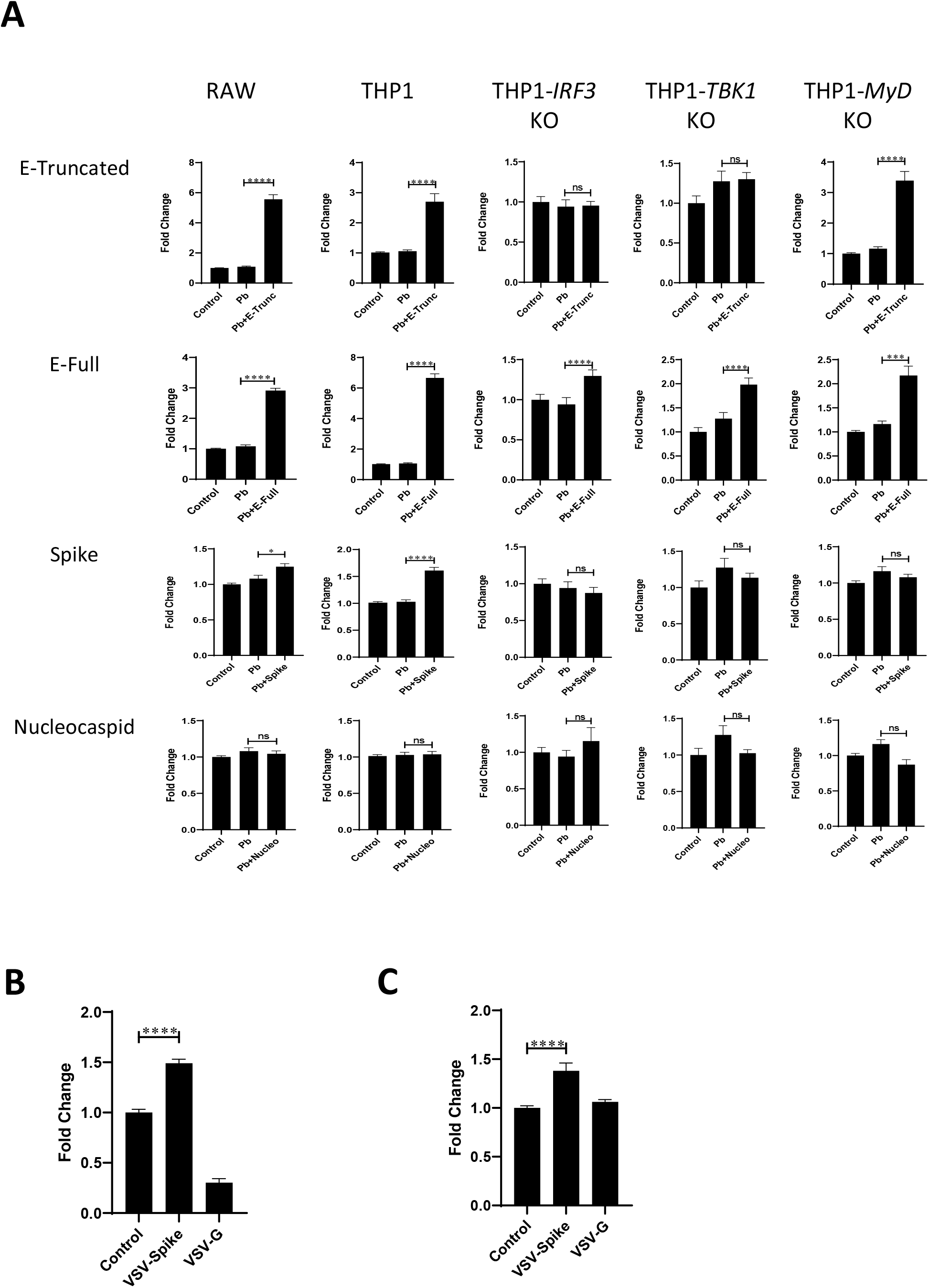
SARS-CoV-2 Proteins Induce IFN Signaling. Fold change in IFN reporter activities in RAW, THP1, THP1-*IRF3*^-/-^, THP1-*TBK1*^-/-^ or THP1-*MyD88* ^-/-^ cells (A) treated with control, polymyxin (Pb) at 10µg/ml and SARS-CoV-2 proteins (E-Trunc, E-Full length, S, N) individually at 2µg/ml with Pb at 10µg/ml for 24 hours (n=2 experiments; 6 biological and 9 technical replicates for RAW with all proteins, n=3 experiments; 9 biological and 6-12 technical replicates for THP1 with all proteins and n=2 experiments; 9 biological and 9 technical replicates for THP1-*IRF3*^-/-^, THP1-*TBK1*^-/-^ or THP1-*MyD88* ^-/-^ cells with all proteins). Fold change in IFN reporter activities in THP1 (B) or RAW (C) cells treated with VSV-SARS-CoV-2-Spike or VSV at MOI:15 for 1 hour and then read at 48 hours. (n=3 experiments; 6 biological and 6 to 17 technical replicates for THP1, n=3 experiments; 6 biological and 6-24 technical replicates for RAW). Graphs depict average with SEM. *p<0.05, ***p<0.001, ****p<0.0001 and ns denotes not statistically significant. Mann-Whitney was used for statistical analysis.

To gain further mechanistic insight into how viral antigens independently activate IFN signaling, we investigated the role of known mediators of IFN induction by pathogen recognition receptors. THP-1 reporter cells deficient in *IRF3* and *TBK1* did not exhibit IFN induction in response to E-Trunc or Spike and had a dramatic decrease in response to E-Full (Fig 2A), demonstrating key dependence on these well described IFN inducing mediators. MyD88 also modulates IFN responses, largely through TLR activation. Indeed, E-Trunc and E-Full showed partial decreases in IFN induction in THP-1-*MyD88* KO cells while the response to S was abolished.

Given that these THP-1 cells are also capable of reporting NFKB induction, we investigated the effect of the above viral antigens in a similar fashion. NFKB induction was also increased in response to viral peptide incubation with the most striking response to E-full (8.5-fold) and least induction to N protein (1.2-fold) (S3A Fig). Overall, *IRF3* and *TBK1* was dispensable for robust NFKB induction while these responses were dependent on *MyD88*, consistent with the well described roles of these mediators in NFKB signaling. Again, VSV-pseudovirus expressing the SARS-CoV-2 S protein induced NFKB signaling, confirming that the S viral peptide expressed in a viral lipid bilayer induces inflammatory signaling in a similar magnitude as solubilized protein (S3B Fig).

Of note, throughout these studies we sought to assure that our findings of IFN and NFKB induction were in response to the viral peptides and not LPS contamination. Using the limulus amoebocyte lysate assay, we found minimal LPS (less than 0.4ng/ml) in our viral protein preparations, consistent with the manufacture’s report. Furthermore, we assessed the responsiveness of our reporter cells lines to similar low doses of LPS (0.1-0.5ng/ml) and found minimal IFN and NFKB responsiveness in THP-1, A549, and RAW cells (S4 Fig A-E). Finally, we performed all IFN and NFKB induction experiments in the presence of 10ug/ml polymyxin B (Pb), a potent LPS neutralizing agent, which inhibited LPS induction of both IFN and NFKB at 1ng/ml (S4F-G Fig).

### Purified SARS-CoV-2 peptides E and S induce lung inflammation and pathology in a mouse model

The critical events that follow an acute pulmonary SARS-CoV-2 infection are injurious viral infection and exuberant immune responses with resultant lung inflammation. Given our findings that E and S can induce similar inflammatory activation pathways in murine cells *in vitro*, we next studied the direct effect of these viral surface antigens *in vivo*. We administered E-Trunc and S intranasally to C57Bl/6J wildtype mice and examined the effect on lung histology three days later. Cross-sections of lungs showed significant organized peribronchial and medium-sized airway pathology in those mice exposed to E-Trunc or S compared to control treated mice (Fig 3A, 3D). Immunostaining for CD45 demonstrated the hematopoietic origin of these inflammatory infiltrates (Fig 3B). Furthermore, animals exposed to E-Trunc and S also showed significant vascular pathology with evidence of vasculitis (Fig 3C), a finding that has been uniquely highlighted in patients with COVID-19 disease (38). These observations demonstrate a striking and direct role of the viral surface proteins in induction of SARS-CoV-2-mediated pathology independent of active viral infection and replication.

**Figure 3.**
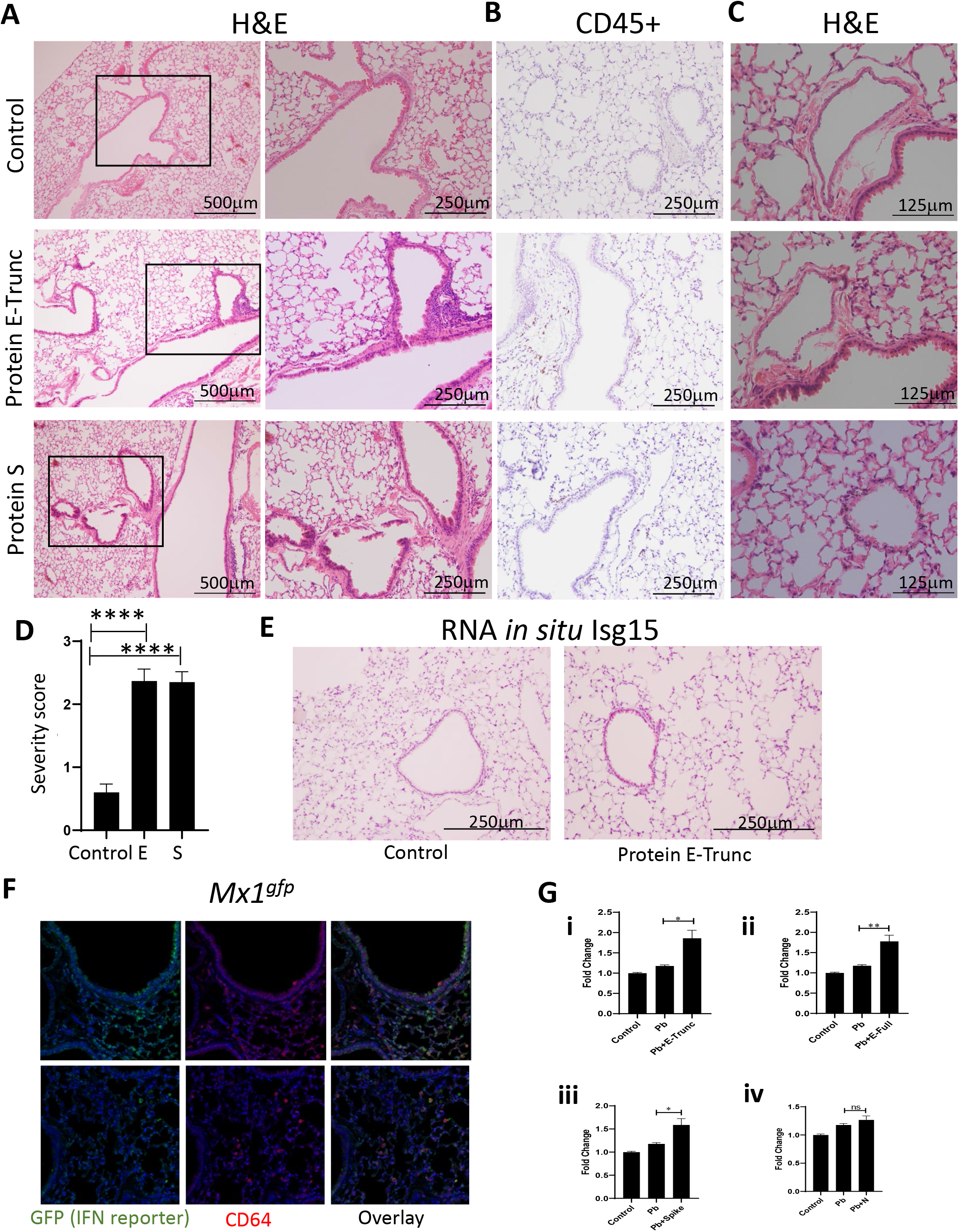
SARS-CoV-2 Protein E Induces Lung Inflammation and Vasculitis in Mice. (A) Representative images of lung cross-sections from mice after intranasal delivery of control or 10µg of E-Trunc and S. H+E stained sections are shown. Boxed areas on the left are magnified to the right. (B) Representative images of the lung cross-sections immunostained for CD45 expression. (C) Representative images of lung cross-sections focused on blood vessels in the above conditions. Scale bars depicted in each picture. (D) Quantification of percent of lobes with inflammatory infiltrates in lungs harvested in the above conditions. (n=3 mice per condition). (E) Representative images of the lung cross-sections stained for Isg15 by RNA *in situ* (n=3 mice per condition). (F) Representative immunofluorescent images of the lung cross-sections immunostained for GFP and CD64 expression 3 days after intranasal delivery of 10ug of E or control (n=2 mice). (G) Fold change in IFN reporter activities in A549 cells with treated with control, Pb at 10µg/ml and SARS-CoV-2 proteins (E-Trunc (i), E-Full length (ii), S (iii), N (iv)) individually at 2µg/ml with Pb at 10µg/ml for 24 hours. (n=2 experiments; 6 biological and 9-21 technical replicates with all proteins). Graphs depict average with SEM. *p<0.05, **p<0.01, ****p<0.0001 and ns denotes not statistically significant. Mann-Whitney used for statistical analysis.

Further investigation revealed IFN activation *in vivo* as E-Trunc-treated animals showed evidence of IFN stimulated gene responses by scattered Isg15 staining by RNA *in situ* as compared to control treated animals (Fig 3E). Notably, Isg15 staining was demonstrated in medium-sized airways as well as scattered peripherally in terminal alveolar spaces. Therefore, we next investigated which cell types respond to viral peptide-mediated IFN signaling *in vivo* using the *Mx1*^*gfp*^ reporter mouse (39). Three days after protein E-Trunc intranasal administration fluorescent GFP+ staining of lung cross-sections demonstrated patchy epithelial cells of medium sized airways and CD64+ cells (monocytes and macrophages) respond to viral peptide induced IFN signaling (Fig 3F).

In light of this airway epithelial IFN-responsiveness, we sought to determine the cell intrinsic induction of inflammatory signaling in A549 reporter cells as an in vitro model of pulmonary epithelium. E-Trunc, E-Full, and S enhanced IFN signaling in reporter A549 pulmonary epithelial cells by approximately 1.5-fold (Fig 3G). N did not induce IFN signaling, and none of these viral peptides increased NFKB induction, although A549 cells are responsive to LPS at low doses (S3A, S4C-D Fig). As A549 cells are known to be susceptible to VSV infection (40, 41), alterations in IFN and NFKB induction in these cells after incubation with VSV-S pseudovirus and VSV-G were limited by visual cytopathic effect (data not shown).

### SARS-CoV-2 E protein-mediated organized inflammation is dependent on type I IFN in pulmonary epithelial cells

To determine the role of type I IFN signaling in SARS-CoV-2 surface protein induction of pulmonary pathology, we rendered the type I IFN signaling pathway defective genetically in the type I IFN receptor (*Ifnar*^*-/-*^) or using an *Ifnar* blocking monoclonal antibody (42) prior to viral peptide treatment. *Ifnar*^*-/-*^ animals or those which received the blocking antibody demonstrated an altered inflammatory infiltrative pattern as demonstrated on lung histological cross-sections compared to wt littermates or isotype treated controls, respectively (Fig 4A-B). These IFN deficient animals had similar scattered immune cells readily apparent in the alveoli spaces whereas the control animals exhibited the previously seen organized infiltrates surrounding medium to large airways.

**Figure 4.**
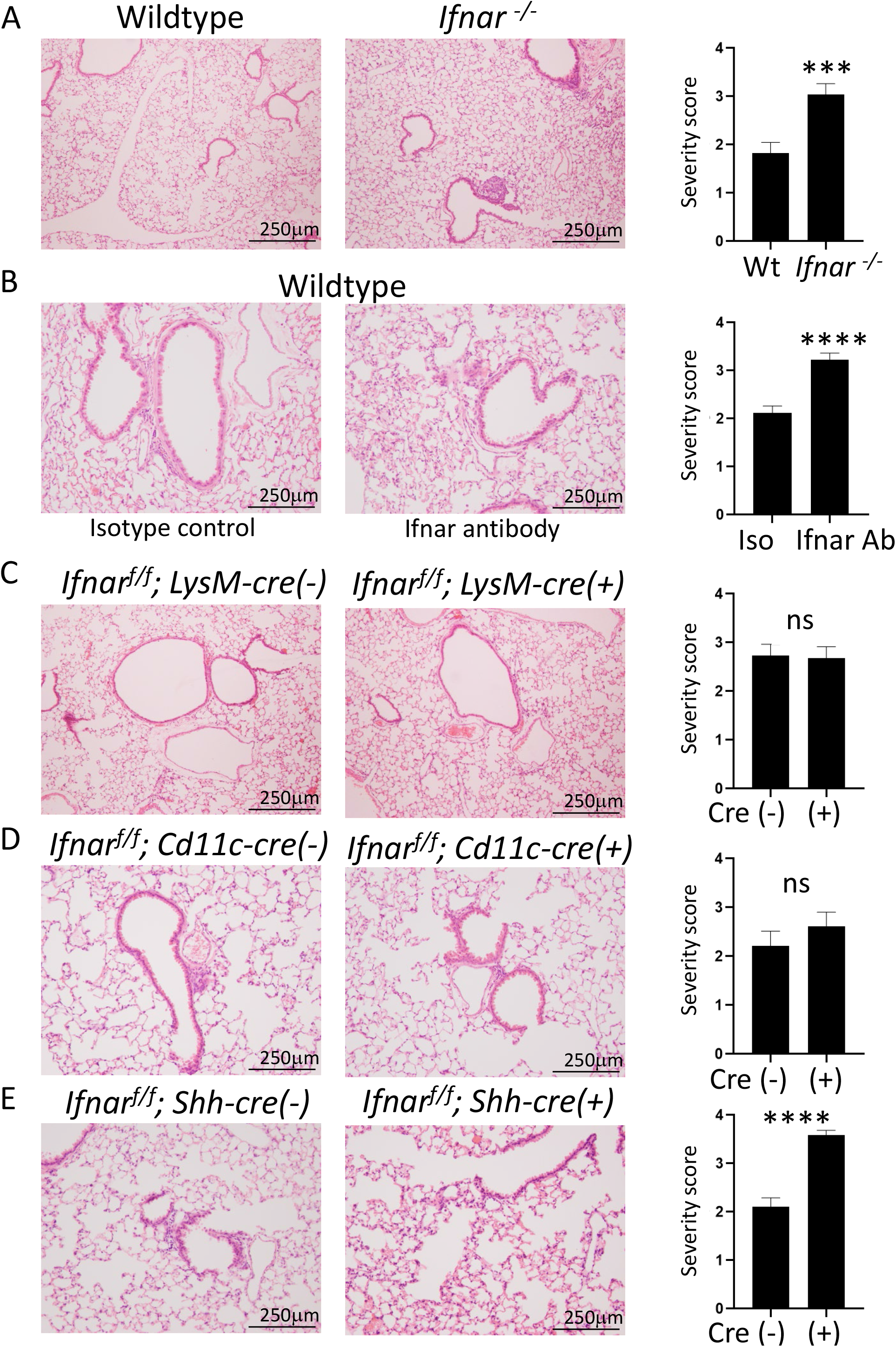
SARS-CoV-2 Protein E Drives Airway Epithelial Cells to Induce IFN-dependent Inflammation. (A) Representative images of lung cross-sections from *Ifnar*^*-/-*^ mice 3 days after intranasal delivery of 10µg of E-Trunc or control. H+E stained sections are shown (n=3 with 4-5 mice total per condition). (B) Representative images of lung cross-sections from mice 3 days after intranasal delivery of 10µg of E-Trunc or control subsequent to IFN-depletion. H+E stained sections are shown (n=2 with 6 mice per condition). (C) Representative images of lung cross-sections from *Ifnar*^*f/f*^;*LysM-Cre(+/*-) mice 3 days after intranasal delivery of 10µg of E-Trunc or control. H+E stained sections are shown (n=3 mice per condition). (D) Representative images of lung cross-sections from *Ifnar*^*f/f*^;*Cd11c-Cre(+/*-) mice 3 days after intranasal delivery of 10µg of E-Trunc or control. H+E stained sections are shown (n=2 mice per condition). (E) Representative images of lung cross-sections from *Ifnar*^*f/f*^;*Shh-Cre(+/*-) mice 3 days after intranasal delivery of 10µg of E-Trunc or control. H+E stained sections are shown (n=3 with 8-9 mice per condition). Scale bars depicted in each picture. Graphs depict average with SEM. ***p<0.001, ****p<0.0001 and ns denotes not statistically significant. Mann-Whitney used for statistical analysis.

Given the dependence on IFN signaling to induce organized inflammation in response to SARS-CoV-2 structural peptides, we utilized a genetic conditional deletion of the type I IFN receptor (*Ifnar*^*f/f*^) (43) in order to determine in which specific cell types IFN signaling determines pulmonary pathology. We targeted the myeloid lineage broadly using LysM-Cre (44) as well as alveolar macrophages and dendritic cells using Cd11c-Cre transgenic mice (45) crossed to *Ifnar*^*f/f*^ mice. SARS-CoV-2 E-Trunc peptide-induced pulmonary pathology was unaffected in *Ifnar*^*f/f*^;*LysM-Cre(+)* and *Ifnar*^*f/f*^;*Cd11c-Cre(+)* compared to their littermate Cre (-) controls (Figure 4C-D). Organized immune infiltrates surrounding medium to large airways was indistinguishable between these groups on lung histological cross-sections. However, mice with type I IFN signaling genetically abolished specifically in pulmonary epithelial cells (46) (*Ifnar*^*f/f*^;*Shh-Cre(+))* exhibited similar pathology to global blockade of IFN signaling in response to E-Trunc (Fig 4E) while the control *Ifnar*^*f/f*^;*Cd11c-Cre(-)* lungs exhibited the afore observed wildtype pathology. These findings implicate the importance of IFN signaling in the pulmonary epithelium as the necessary driver of organized medium to large airway inflammation in response to SARS-CoV-2 surface antigens.

## Discussion

The novel coronavirus SARS-CoV-2 that has caused the COVID-19 global health crisis necessitates a thorough investigation of the host immune response in order to develop effective therapeutic strategies. The innate immune system is the first line of defense that is critical for viral pathogen clearance, and at the same time, it is also implicated in the pathogenesis of many viral disease processes (47). Studies have shown that a hyperinflammatory state and a dysregulated immune response may underlie COVID-19 pathogenesis (1, 2, 7-9, 24-27, 48). COVID-19 patients experience a characteristic “cytokine storm” with sharply high levels of proinflammatory mediators, which is directly proportional to viral load and severity of illness (1, 2, 7, 25, 26, 29-34, 48-50).

While viral nucleic acid-sensing is the predominantly accepted mechanism for virus detection by pathogen-recognition receptors, viral surface proteins may also directly activate the innate immune system independent of virus uncoating and replication. This knowledge is crucial to understand the initiation of the inflammatory response and the mechanism of viral engagement of the immune system. In addition, this mechanism is important to consider given its implications for non-infected cells to induce an immune response as recognition of viral antigens may occur independently of viral uncoating and replication, and thus may not be restricted to cells or tissues permissive to infection.

Hence, we evaluated the ability of isolated SARS-CoV-2 structural antigens to activate IFN signaling, a key innate immune pathway that bridges to adaptive immune responses. Our findings demonstrate that the SARS-CoV-2 surface peptides E and S independently activate IFN signaling in both immune and epithelial cells. We show that these viral antigens individually alter the expression of key chemokines and cytokines, including many regulated by IFN, in both human and murine cell lines. Distinctly, the truncated E peptide enhanced the levels of human CCL5, IP-10, CXCL1, IL-8, CCL2, and MIP-1α, which are associated with neutrophil and monocyte recruitment. These findings are essential in the light of *in vivo* infection, as COVID-19 patients often have a high ratio of neutrophils to lymphocytes (30). In murine cells, the E protein led to increased levels of TNF-α and G-CSF, which are also markedly increased in human SARs-CoV-2 infection (1, 21). Furthermore, we demonstrate that *in vivo* delivery of these peptides, particularly E-Trunc, to mice induces peribronchial and medium-sized airway inflammation and vasculitis, which are recapitulated in human disease specimens (38, 51). This inflammatory recruitment is dependent on IFN signaling in epithelial cells as specific genetic IFN signaling deficiency in pulmonary epithelial cells abolished organized inflammation, although alveolar inflammatory infiltrates persisted. These findings indicate that the pulmonary epithelium can induce IFN signaling and localized inflammation in response to SARS-CoV-2 viral surface protein recognition. Similarly, the SARS-CoV E protein induced severe lung pathology including profuse hemorrhage and cellular infiltration with elevation of cytokines (52). The significance to disease pathogenesis of these inflammatory responses with distinct pathological patterns warrants further assessment in genetically modifiable host-pathogen and SARS-CoV-2 host-susceptible model systems.

While the S protein is responsible for cell entry via ACE2 and is the focus of numerous therapeutic strategies, the E protein of SARS-CoV-2 is understudied although recent evidence points to its potential as an ion channel (53). Prior work in other coronaviruses has demonstrated that E protein is indispensable for viral morphogenesis and tropism as well as enhances inflammasome activation (14-18); our work further points to its crucial role on innate immune activation and function. Given that E is highly conserved with SARS-CoV (9, 54) further study is necessary as E may be a potent target for therapeutic strategies with broader applications, including anticipated emerging coronaviruses. Our observations also highlight the importance of the direct effect of coronavirus surface proteins and will usher investigation of other viral surface proteins as determinants of the host-pathogen interaction.

Finally, this work has broad implications for the pathogen-host immune response; we show that activation of innate immune signaling pathways independent of viral nucleic acid detection by pathogen recognition receptors engage host immunity similarly to complete infectious virus. Understanding the immune response to independent viral structural proteins is an important step forward in deciphering the interaction of this novel virus, as well as other clinically relevant viruses, with host immunity.

## Materials and Methods

### Mice

All mice were originally obtained from Jackson Laboratories (Bar Harbor, Maine, USA) and subsequently maintained at Washington University under specific pathogen-fee conditions and bred in-house. Adult (8-16 week-old male and female) mice were anesthetized with isoflurane and intranasally administered 10µg of protein E, S, or water in 50µl total volume (25 µl per nare). Mice were sacrificed on day 3 post-administration, and lung specimens isolated and evaluated by histology. For the IFN-depleting experiments, mice were injected intraperitoneally with 2mg antibody in 500µl volume (anti-Ifnar or isotype control) 6 days prior and 0.5mg antibody in 500ul volume 2 days prior to intranasal administration of protein E.

### Cell lines

The cell lines A549-Dual (adenocarcinoma human alveolar basal epithelial cells) and RAW-Lucia ISG (RAW-mouse macrophages) (InvivoGen, San Diego, CA, USA) were cultured in DMEM (Sigma-Aldrich, St Louis, Mo, USA). The growth media was supplemented with 10% fetal bovine serum (FBS) (Sigma-Aldrich), 1% (v/v) of penicillin/streptomycin, 100 µg/ml Normocin/Zeocin (InvivoGen). The A459 cells were also supplemented with 100 µg/ml Blasticidin. The THP1-Dual and THP1-Dual KO (IRF3/TBK1/MyD) cells (human lung monocytes) were cultured in RPMI 1640 (Sigma-Aldrich) medium supplemented with 10% heat-inactivated FBS, 2mM L-glutamine, 25mM HEPES (Sigma-Aldrich), 1% (v/v) of penicillin/streptomycin and 100 µg/ml of Normocin/Zeocin/Blasticidin (InvivoGen). The test media for A549 and THP-1 cells excluded Zeocin and Blasticidin from their respective growth media.

### Reporter cell assays

A549-Dual, THP1-Dual, THP1-Dual KO and RAW-Lucia ISG cells stably express an interferon regulatory factor-inducible Lucia luciferase reporter construct. Cells were seeded at 1×10^6^ or 1×10^5^ cells per well in a 6-well or 96-well plate, respectively. Cells were then incubated with S, E, or N protein at 2 µg/mL or equal volume of water as control, and after 24 hours, culture supernatant was collected to measure luciferase. QUANTI-Luc was used to detect the level of luciferase by adding to culture supernatant and reading immediately with a plate reader (Infinite M200 Pro, TECAN Life Sciences, Switzerland) at a 0.1 second reading time. QUANTI-Blue was used to detect the level of SEAP (secreted embryonic alkaline phosphatase) by adding to culture supernatant and incubating for 1 hour and reading with a plate reader (Infinite M200 Pro, TECAN Life Sciences, Switzerland) at 650 nm.

### Chemokine and cytokine analyses

Chemokine and cytokine protein quantification were performed using Proteome Human and Mouse Cytokine Array kits (R&D Systems, San Diego, CA) as per the manufacturer’s instructions. Dot arrays were quantified for pixel density with ImageJ.

### Lung tissue preparation for histology

Lungs were inflated with formalin at the time of sacrifice and harvested into formalin containing conical tubes. The tissue was serially washed with PBS (Sigma-Aldrich), 30% ethanol, and 50% ethanol 48 hours after harvesting and stored in 70% ethanol until processed for paraffin embedding, sectioning, and staining.

### Software

ZEN 3.1 blue edition was used to visualize and image immunofluorescence staining of lung sections.

### Statistics

GraphPad Prism (San Diego, CA, USA) version 7.02 software was used to perform all statistical analyses as described.

### Ethics Statement

All animal protocols used in this study were approved by the Washington University’s Animal Studies Committee (19-0768), which approved these methods. Humane sacrifice of animals occurred with isoflurane administration and cervical dislocation.

## Acknowledgements

Pseudoviruses VSV-G and VSV-Spike were kindly supplied by Dr. Sean Whelan, Washington University in St. Louis.

---

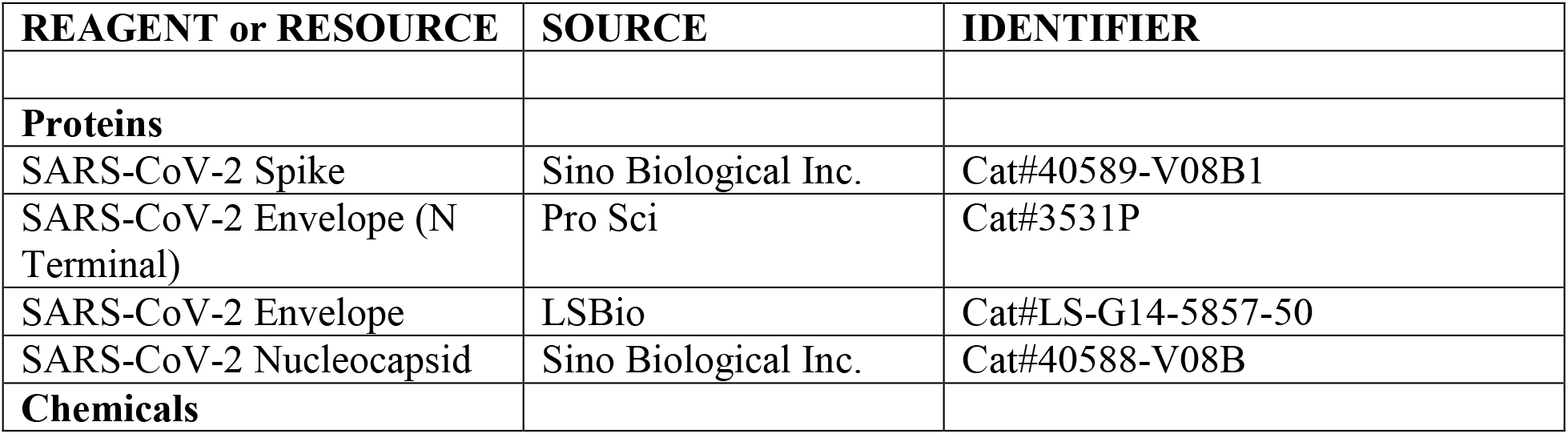
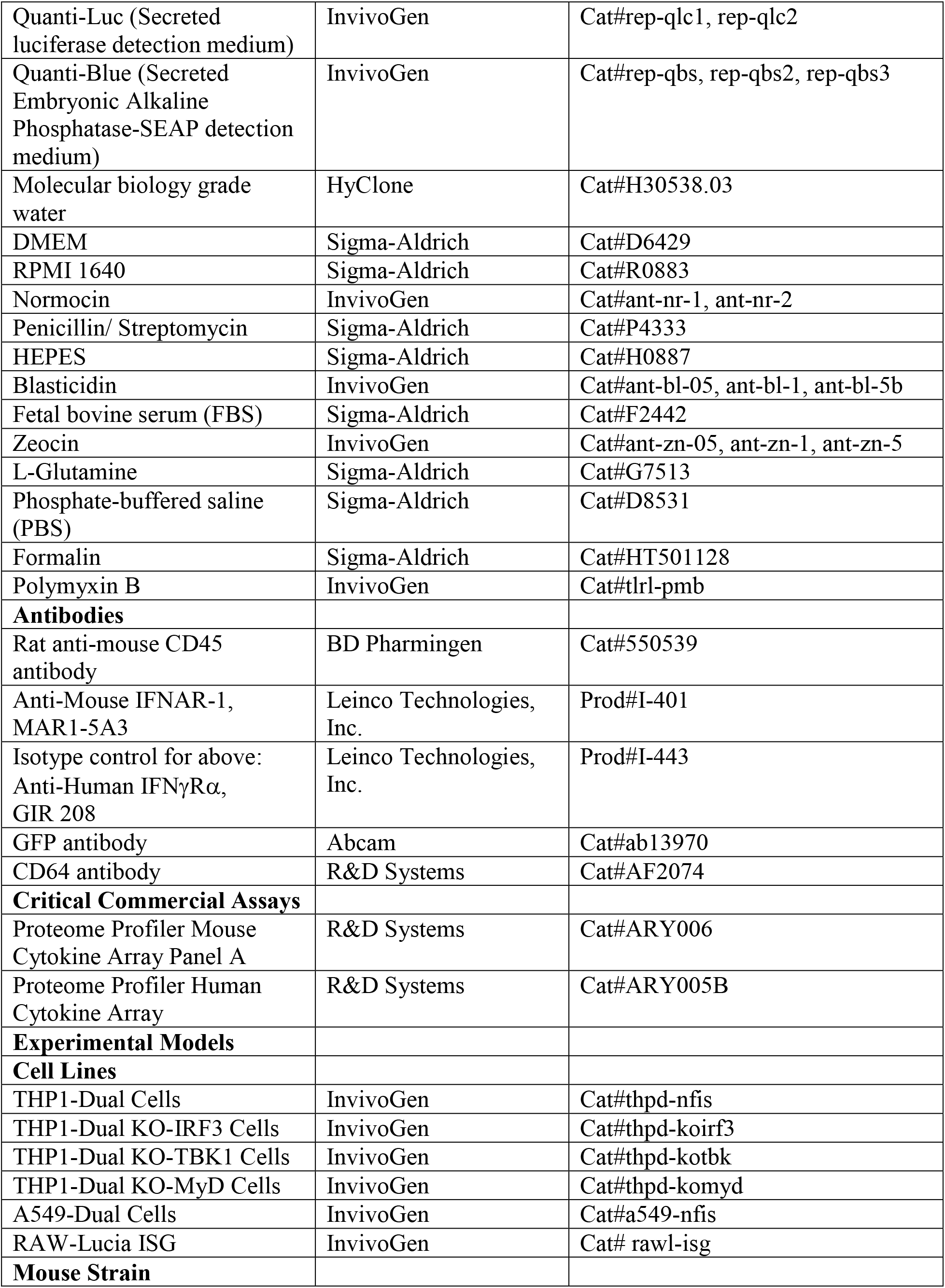
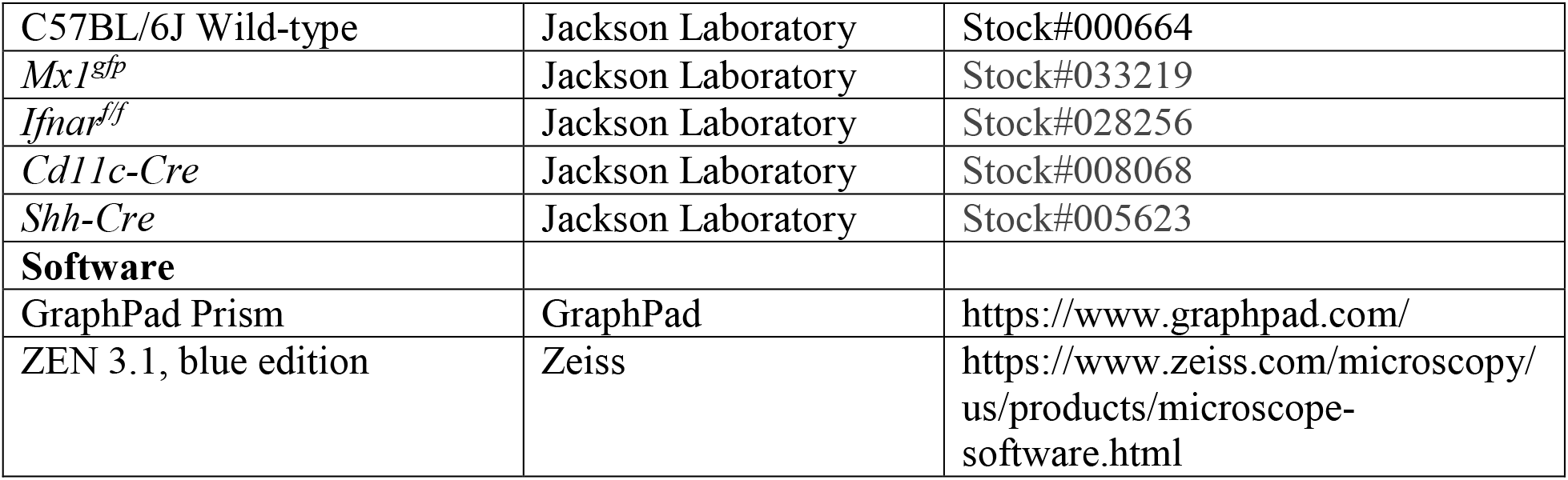

## Lead Contact

Further information and requests for resources and reagents may be directed to and will be fulfilled by the corresponding author, Ashley L. Steed.

