## Supplement for "Surface proteins of SARS-CoV-2 drive airway epithelial cells to induce interferon-dependent inflammation"

### Supporting information

**S1 Fig. SARS-CoV-2 Proteins E and S Elevates the Expression of Proinflammatory Chemokines and Cytokines in Murine and Human Cells.** Detection of chemokines and cytokines in the culture supernatant of RAW (A) and THP-1 (B) cells incubated with control, E-Trunc or S at 2μg/ml for 24 hours. The graphs show measurements of pixel density in the dot arrays under control and experimental conditions (n=2 biological samples for each condition with 2 technical replicates of each sample). Graphs depict average with SEM. *p< or =0.05 and ns denotes not statistically significant. Mann-Whitney used for statistical analysis.

**S2 Fig. SARS-CoV-2 Proteins E and S Elevate the Expression of ISGs.** Enhanced expression of CCL5 (A) and TNF-α (B) detected from THP-1 cells incubated with control, E-Trunc or S at 2μg/ml for 3 hours (n=2 experiments; 4 biological and 4 technical replicates of each sample). Graphs depict average with SEM. *p< or =0.05 and ns denotes not statistically significant. Mann-Whitney used for statistical analysis.

**S3 Fig. SARS-CoV-2 Structural Proteins Induce NF-kB Signaling.** Fold change in NF-kB reporter activities in THP1, THP1-*IRF3*$^{-/-}$, THP1-*TBK1*$^{-/-}$, THP1-*MyD8*8$^{-/-}$ cells or A549 (A) treated with control, Pb at 10μg/ml and SARS-CoV-2 proteins (E-Trunc, E-Full length, S, N) individually at 2μg/ml with Pb at 10μg/ml for 24 hours (n=3 experiments, 9 biological and 6-12 technical replicates for THP1 with all proteins; n=2 experiments, 9 biological and 9 technical replicates for THP1-*IRF3*$^{-/-}$, THP1-*TBK1*$^{-/-}$, THP1-*MyD8*8$^{-/-}$ cells with all proteins; n=2 experiments, 6 biological and 8-20 technical replicates for A549 with all proteins). Fold change in NF-kB reporter activities in THP1 (B) cells treated with VSV-SARS-CoV-2-Spike or VSV at

MOI:15 for 1 hour and then read at 48 hours. (n=3 experiments; 6 biological and 6 to 12 technical replicates for THP1). Graphs depict average with SEM. *$p<0.05$, **$p<0.01$, ****$p<0.0001$ and ns denotes not statistically significant. Mann-Whitney used for statistical analysis.

**S4 Fig. Reporter Cell Lines Responsiveness to LPS.** Fold change in IFN (A) and NF-kB (B) reporter activities in THP-1 cells in response to low-dose LPS. Fold change in IFN (C) and NF-kB (D) reporter activities in A549 cells in response to low-dose LPS. Fold change in IFN (E) reporter activity in RAW cells in response to low-dose LPS. (n=2 experiments; 12 biological and 24 technical replicates). Fold change in IFN (F) and NF-kB (G) reporter activities in THP-1 cells in response to 1ng/ml LPS in the presence of increasing concentrations of Pb.

**S5 Fig. Pulmonary Inflammatory Infiltrate Score.** Score 0: no appreciable inflammation surrounding airways and in the alveolar spaces. Score 1: minimal inflammatory infiltration limited to central airways, Score 2: moderate inflammatory infiltration surrounding central airways with some appreciable peripheral inflammation, Score 3: moderate inflammatory infiltration surrounding central airways with moderate peripheral inflammation. Score 4: moderate inflammatory infiltration surrounding central airways with widespread peripheral inflammation.

Supplemental Figure 1

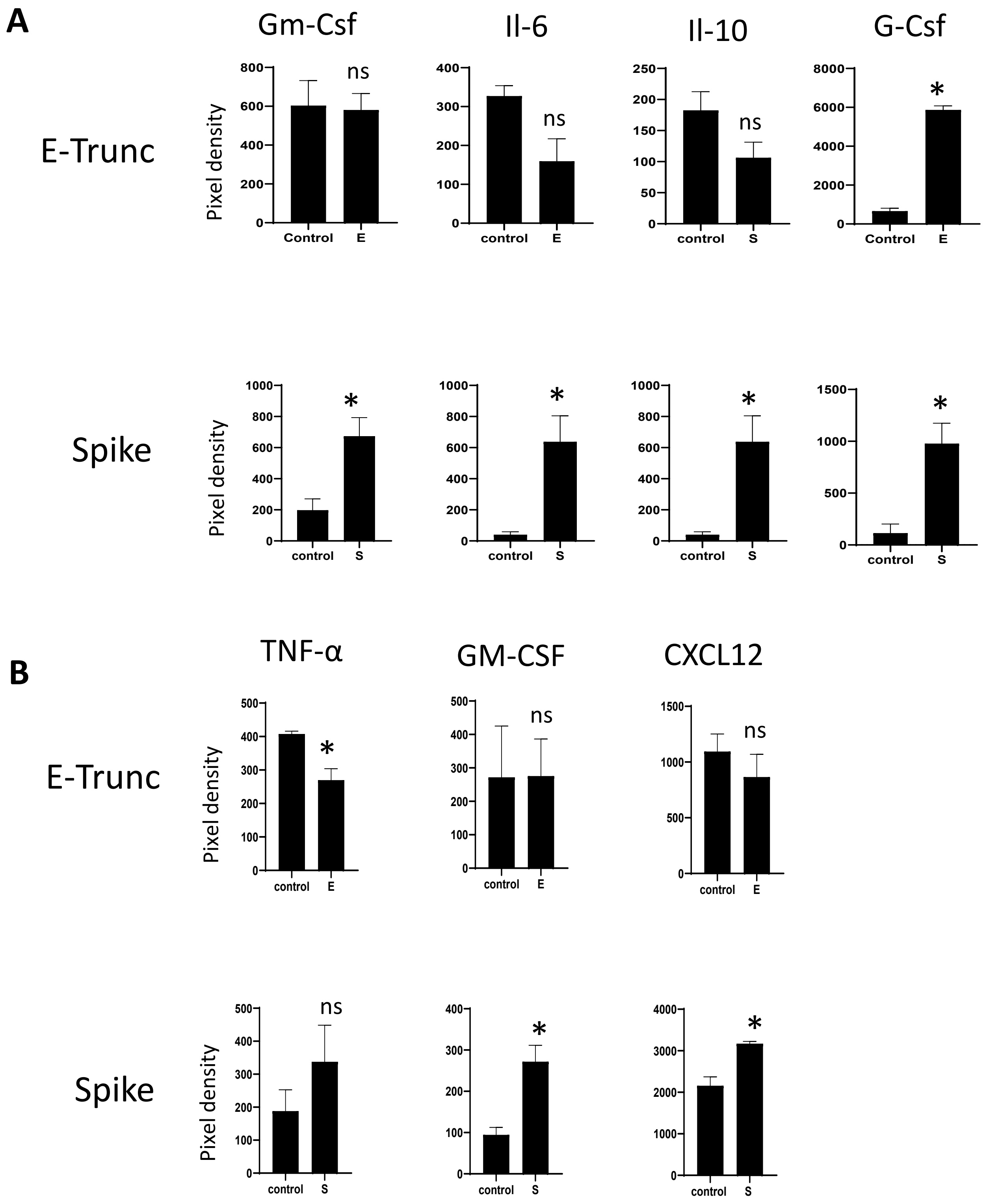

Supplemental Figure 2

A

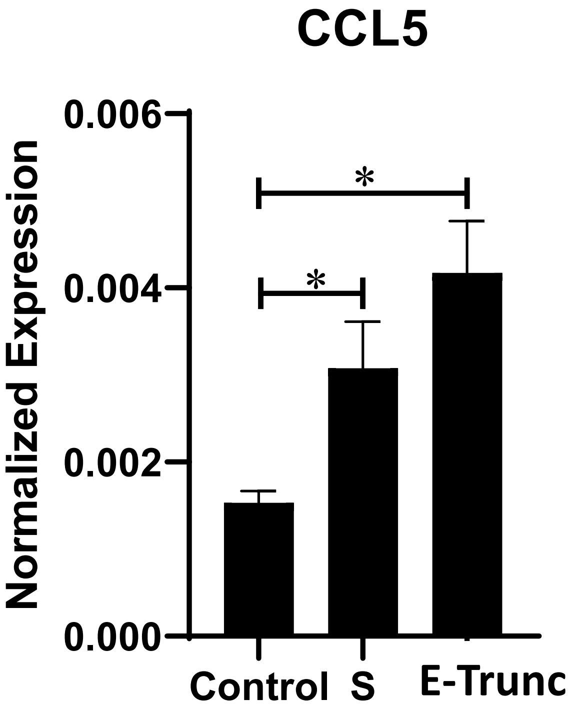


B

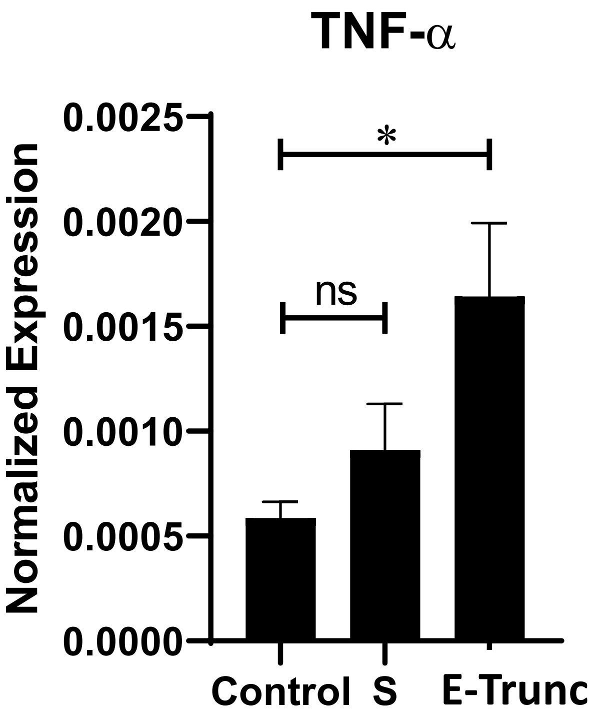

Supplemental Figure 3

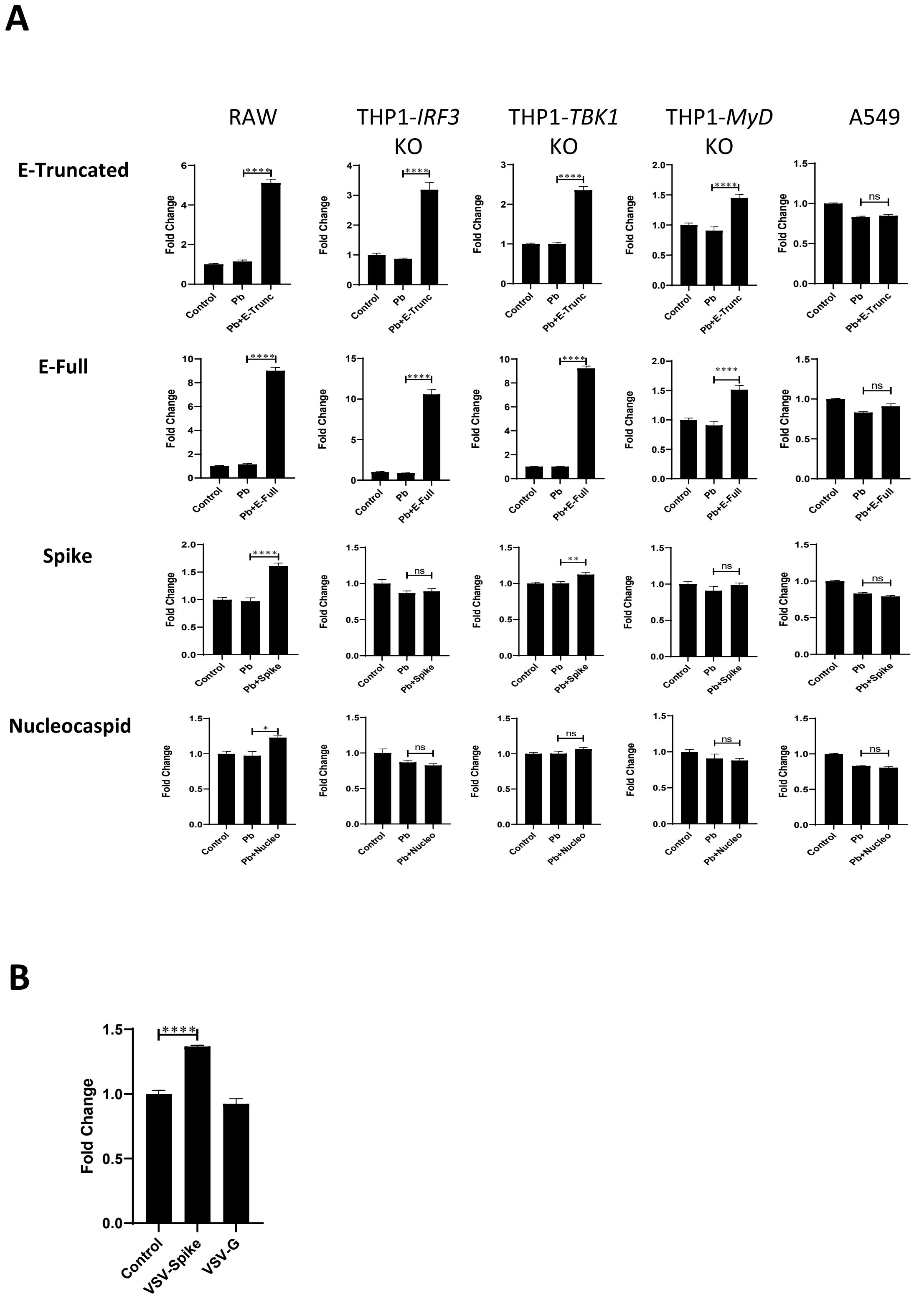

Supplemental Figure 4

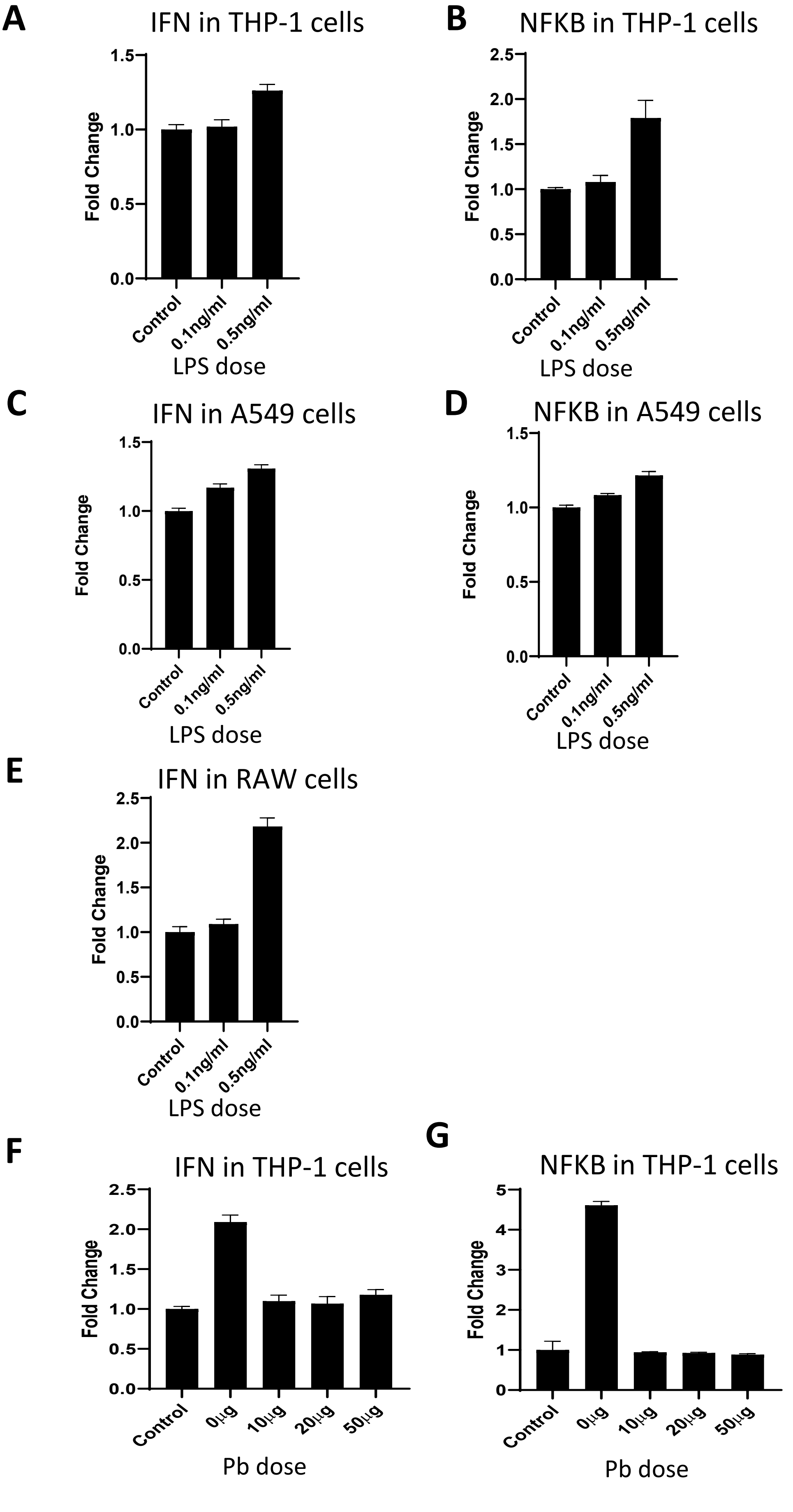

### Supplemental Figure 5

Score 0

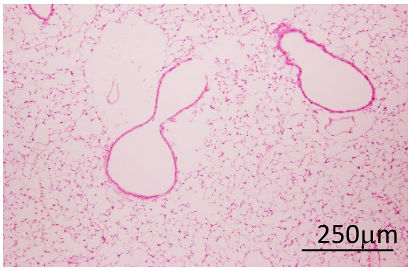


Score 1

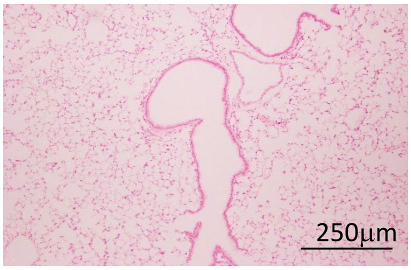


Score 2

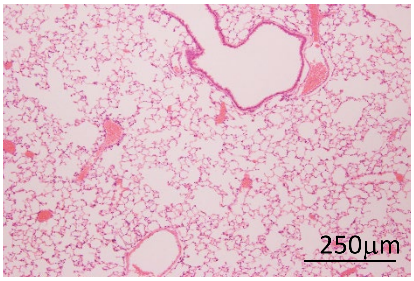


Score 3

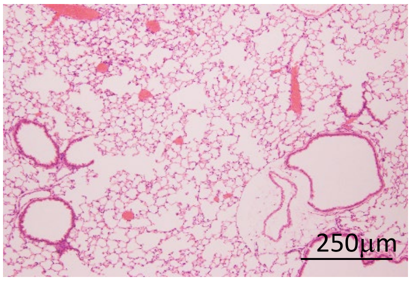


Score 4

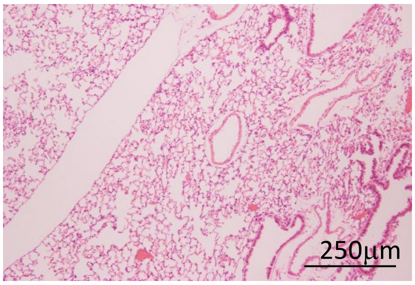
